# Parvalbumin loss following chronic sub-anesthetic NMDA antagonist treatment is age-dependent in the hippocampus: Implications for modeling NMDA hypofunction

**DOI:** 10.1101/399188

**Authors:** Jennifer A. Honeycutt, James J. Chrobak

## Abstract

A marked decrease in parvalbumin (PV), a calcium-binding protein specific to a subset of GABAergic neurons, is a consistent finding in postmortem schizophrenic brain tissue. This reduction is selective to PV and is regionally specific: occurring primarily in the prefrontal cortex and hippocampus (HPC) of patients. Rodent models of NMDA receptor hypofunction utilizing NMDA antagonist treatments – e.g. ketamine (KET) – show schizophrenia-like cognitive and behavioral impairments with parallel changes in PV. While decreased PV is considered a hallmark of neuropathology in schizophrenia, previous work elucidating the effects of KET administration on PV are contradictory, with findings suggesting decreased, increased, or no change in PV expression. Upon close examination of the procedures used across studies, there are two primary inconsistencies, including: 1) the age of animals used; and 2) the timeline of post-treatment tissue collection. To better understand whether these key differences impact observed PV changes, the present study investigated the impact of age and time of sacrifice on chronic KET-induced PV changes in the neocortex and HPC. Our findings suggest an effect of age, but not sacrifice timeline, on PV cell count following 14 days of sub-anesthetic KET treatment. We provide evidence that 1-month-old rats exhibit significant KET-induced HPC PV *decreases*, while adult rats show a modest *increase* in HPC PV following chronic KET. Taken together, we propose that PV is a dynamic marker, and that changes in cell counts - and their interpretation - following NDMA antagonist treatment should be considered in the context of age.

## Introduction

Investigations into the neuropathology of schizophrenia in patient postmortem brain tissue have revealed abnormalities in neuroanatomical structure and changes in biological markers of distinct neural populations (Eyles et al., 2002). These observed abnormalities have given us a glimpse of putative mechanisms that may underlie the pervasive cognitive and behavioral changes characteristic of schizophrenia. Furthermore, these findings have allowed for the advancement of rodent modeling techniques, which have become integral to our ability to systematically study possible factors contributing to neural and behavioral pathological phenotypes. Specifically, postmortem findings reliably point to a selective decrease in the calcium-binding protein parvalbumin (PV) exclusively within the prefrontal cortex (PFC; Beasley and Reynolds, 1997) and hippocampus (HPC; Zhang and Reynolds, 2002). In the rodent literature, models employing N-methyl-D-aspartate (NMDA) antagonist treatments (e.g. ketamine (KET); MK-801), as a means of inducing a state of NMDA receptor hypofunction, are widely accepted and used in large part for their ability to replicate cognitive and behavioral dysfunction seen in schizophrenia (Meltzer et al., 2011; Coyle et al., 2012; Meltzer et al., 2013). Importantly, NMDA antagonist treatments in rodents have been shown to replicate the robust and specific decrease in PV expression seen in schizophrenic patients within the HPC and PFC (Reynolds et al., 2004; Adell et al., 2012; Gonzalez-Burgos and Lewis, 2012). However, despite general agreement that decreased PV expression (via protein levels and/or cell counts) following pharmacologically-induced NMDA hypofunction is a valid model of schizophrenic neuropathology, there exists a growing number of studies presenting conflicting findings, calling into question the aptness of utilizing PV as a marker of pathology across development.

Within the literature, discrepancies regarding the outcomes of NMDA antagonism on neuropathology, particularly on PV, are apparent with some groups indicating decreases in PV following treatment (e.g., Keilhoff et al., 2004a; Keilhoff et al., 2004b; Behrens et al., 2007; Braun et al., 2007; Wang et al., 2008; Zhang et al., 2008; Rómon et al., 2011; Kittelberger et al. 2012), while others report no change following treatment (e.g., Rujescu et al., 2005; Benneyworth et al., 2011). These conflicting findings raise questions regarding the methodological preparations employed, as well as the translational validity of the findings. Interestingly, upon examination of the methods used across various studies, critical differences in methodology exist including: 1) NMDA antagonist treatment regimen; 2) the age of the subjects; 3) whether or not subjects are behaviorally naïve; and 4) the timeline of brain tissue collection following the cessation of treatment and/or behavioral assays (for a more comprehensive list of previous findings with detailed methods (i.e. age, treatment regimen, sacrifice timeline, etc.) see Table 1). Indeed, the lack of continuity and comprehensive rationale in the age of subjects used across studies points to an underlying assumption of stability in PV phenotype over the lifespan, which may prove problematic when trying to translate findings to additional populations of interest.

**Table 1.** Previous work detailing changes in PV pathology following NMDA antagonist treatment.

| Study | Species | Age | Drug | Treatment Regimen | Timeline: Post-Cessation | Quant. Method | PV Results |  |
| --- | --- | --- | --- | --- | --- | --- | --- | --- |
|  |  |  |  |  |  |  | HPC | PFC |
| Abdul-Monim et al., 2007 | Rat (f) | Not specified (approx. >5mo) | PCP | 2mg/kg (2x/d) x 7 days | 42 days | IHC | ↓(CA3, DG) | ↓(M1)<br>↑(M2; CG) |
| Behrens et al., 2007 | Mouse (m) | Not specified | KET | 30mg/kg (IP) x 2 days | 18 hours | IHC | -- | ↓ |
| Behrens et al., 2008 | Mouse (m) | 12-15 weeks | KET | 30mg/kg (IP) x 1 or 2 days | 1, 3, or 10 days | IHC | -- | n.c. (1d)<br>↓ (2d) |
| Benneyworth et al., 2011 | Rat (m) | Not specified | KET | 30mg/kg (IP) x 5 days | 72 hours | IHC; WB | n.c. | n.c. |
|  | Mouse (m) | 9-13 weeks | KET or PCP | 30mg/kg (IP) x 5 days<br>6mg/kg (IP) x 5 days | 72 hours | IHC; WB | n.c. | n.c. |
| Braun et al., 2007 | Rat (m) | Approx. 35 days | MK-801 | .02mg/kg (IP) x 21 days | 24 hours | IHC | ↓(CA1, DG) | n.c. |
| Jeevakumar et al., 2015 | Mouse (m) | 1-17 weeks | KET | 30mg/kg (SC) on PD7, 9, 11 | 16 weeks | IHC | -- | ↓ |
| Jenkins et al., 2010 | Rat (m) | Not specified | PCP | 2 or 5mg/kg (IP) x 7 days | 42 days | IHC | ↓(CA1, CA3) | -- |
| Keilhoff et al., 2004 | Rat (m) | 8 weeks | KET | 30mg/kg (IP) x 5 days | 14 days | IHC | ↓(CA1, CA3, DG) | -- |
| Kittelberger et al., 2012 | Rat (m) | Adult (age not specified) | KET | 30mg/kg (IP) x 5 days | 2-4 weeks | IHC | ↓(CA1, CA3) | -- |
| Koh et al., 2016 | Mouse (m) | 1-6 months | KET | 16mg/kg (SC) x 1 month (3x/week) | 3-5 months | IHC | ↓(CA1) | n.c. |
| Rujescu et al., 2005 | Rat (m) | 32-40 days | MK-801 | .02mg/kg (IP) x 14 days | 24 hours | IHC | ↓(HPC total)<br>n.c. (CA1, CA3, DG) | -- |
| Schobel et al., 2013 | Mouse (m) | 65-75 days | KET | 16mg/kg (IP) x 1 month (3x/wk) | 48 hours | IHC | ↓(CA fields) | -- |

Previous work evaluating the impact of acute and/or chronic NMDA antagonism varies widely in methodologies utilized, particularly with regard to length and dose of treatment, age of subjects, amount of time between treatment-cessation and tissue collection. These discrepancies likely contribute to variability between studies and research groups.
Abbreviations: ↓, decrease from CON; ↑, increase from CON; n.c., no change from CON; f, female; m, male; PD, Postnatal Day; IHC, Immunohistochemistry; WB, Western Blot; M1, Primary Motor Cortex; M2, Secondary Motor Cortex; CG, Cingulate Cortex; IP, Intraperitoneal Injection; SC, Subcutaneous Injection

Findings indicating developmental changes in PV from young to adult rodents provide a compelling argument for considering age as a factor. Specifically, our group and others have presented evidence of decreased PV within the HPC in adult compared to young rodents (de Jong et al., 1996; Honeycutt et al., 2016). These findings indicate a distinct developmental trajectory of PV phenotype which likely influences the efficacy and/or outcomes of experimental treatments based on subject age. In addition to age mediating PV outcomes across studies, there is evidence that behavioral and cognitive training/testing may also serve to alter PV across multiple brain regions (e.g., Gomes da Silva et al., 2010; Urakawa et al., 2013). Thus, it is necessary to determine the baseline effect of pharmacological models in the absence of behavioral manipulations to obtain a clear understanding of the effect of the KET model in isolation. In addition to these possible mediating factors, time of tissue collection – with respect to cessation of treatment – may be a significant mediator of PV findings, as work by Behrens and colleagues (2008) suggests that PV phenotype may recover as a function of time post treatment. These factors, which have been largely overlooked as a significant driving influence in PV changes following NMDA antagonist treatment, may provide key insights into the dynamic nature of PV and elucidate its relationship to pathology in schizophrenia.

A more thorough understanding of the normative relationship between PV and these possible mitigating factors is warranted, thus the present work sought to investigate: 1) age-dependent changes in PV across development in behaviorally naïve rats; 2) the neural consequences of chronic KET administration on PV as a function of age; and 3) the role of tissue collection timeline post KET cessation on PV cell counts in key brain regions known to be selectively impacted in schizophrenia. Here, we provide evidence suggesting a differential effect of chronic KET administration as a function of age, such that 1-month-old rats show a significant decrease in PV, whereas 6-month-old rats show a pattern of increased PV within the HPC. To our knowledge, this is the first report detailing a distinct and opposite pattern of PV+ cell counts within the HPC following chronic KET treatment in an age-dependent manner. Taken together, these findings provide important insight into the need to carefully consider stage of development in rodent models of pathology.

## Methods

### Subjects and drug administration

Subjects were male Sprague-Dawley rats (*n*=30) sourced from Charles River Laboratories (Wilmington, MA) and aged either 1 (*n*=15) or 6 months (*n*=15). Prior to pharmacological manipulations, rats were individually housed in polycarbonate caging with free access to food and water and were allowed 1 week to acclimate to the colony room. All rats were housed in a temperature and humidity-controlled vivarium with a 12-hour light/dark cycle (lights on at 08:00h) and exposed to normal weekly husbandry procedures but were otherwise behaviorally naïve. Subsets of rats from each age group were assigned to one of 3 treatment groups: 1) chronic KET regimen with brain tissue collection immediately following treatment cessation (KET-I; 30 mg/kg IP; *n*=5 per age group); 2) chronic KET regimen with tissue collection delayed 10 days post treatment cessation (KET-D; 30 mg/kg IP; *n*=5 per age group); or 3) comparable control with IP injection of physiological saline (CON; *n*=5 per age group). Chronic KET administration was sub-anesthetic and lasted for 14 consecutive days (see Figure 1A for experimental timeline). Importantly, the chronic KET regimen did not impact the overall health of the animals, as evidenced by no significant weight changes between groups across treatment days (data not shown). The KET dose of 30 mg/kg was chosen based on the frequency with which it is used in the literature by many research groups as a sub-anesthetic acute and/or chronic treatment model of NMDA hypofunction (see Table 1). All housing and experimental procedures were performed in accordance with, and approved by, the University of Connecticut Institutional Animal Care and Use Committee. See Figure 1A for a methodological timeline.

**Figure 1.**
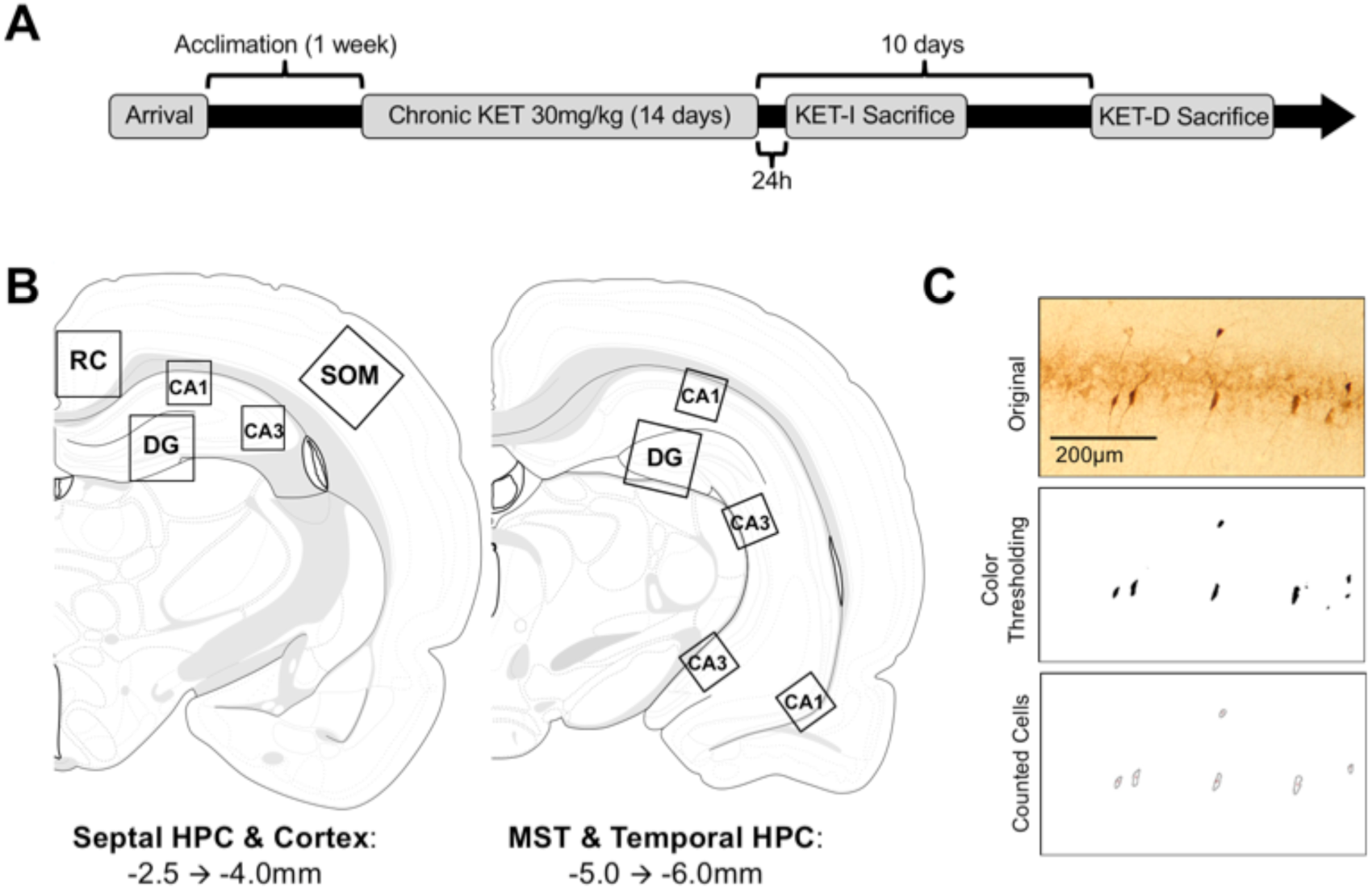
Experimental Methods: Methodological timeline of experiment (**A**). After a one-week acclimation rats began chronic treatment (either 30mg/kg KET or saline CON, IP) for 14 consecutive days. Rats were then sacrificed either 24 hours (KET-I) or 10 days (KET-D) following treatment cessation. Sampling regions (**B**) included regions encompassing the retrosplenial cortex (RC), Somatosensory Barrel Fields (SOM), and HPC sub-regions (CA1, CA3, DG) across the septotemporal axis. An example of PV+ cell quantification within HPC CA1 can be seen in (**C**). The top panel shows an original photomicrograph with darkly stained PV+ cells. The center panel shows the same photomicrograph thresholded based on color density to isolate the most darkly stained cells. A macro as run to quantify the number of PV+ cells while excluding non-cell artifacts, with the output indicating counted cells shown in the bottom panel. All anatomical measurements are relative to Bregma. Atlas adapted from Swanson (2004).

### Tissue Collection & Immunohistochemistry

Following treatment cessation and experimental timeline, rats were anesthetized with Euthasol (pentobarbital solution; 0.5-1.0mL delivered IP) and transcardially perfused with ice-cold 0.9% physiological saline solution immediately followed by ice-cold 3.7% paraformaldehyde solution. Brains were removed and individually stored in 3.7% paraformaldehyde solution for 1 week. Prior to sectioning, brains were transferred into a 30% sucrose solution for cryoprotection. All brains were coronally sliced into 60μm serial sections on a cryostat and stored in phosphate-buffered saline (PBS)-containing well plates prior to immunohistochemical procedures.

One series of tissue from each brain was selected to examine PV immunoreactivity (distance per analyzed slice approx. 360μm), with sections encompassing tissue approximately −2.5mm through −6mm relative to Bregma to include the septotemporal extent of the HPC, as well as adjacent cortical regions (retrosplenial and somatosensory cortices; Paxinos and Watson, 1997). Free-floating sections were initially blocked in 5% normal goat serum (NGS; Jackson Immuno), 0.1% triton-X, and PBS solution for 1 hour, followed by 3 five-minute washes in PBS. Sections were transferred to primary antibody solution (1:16,000 rabbit anti-parvalbumin polyclonal (ABCAM), 0.1% triton-X, and PBS) for a 1-hour incubation at room temperature. Sections were rinsed in PBS and transferred to secondary solution (horseradish peroxidase (HRP) labeled polymer anti-rabbit; DAKO) for 1 hour at room temperature. Following a final set of washes in PBS, sections were transferred into a diaminobenzidine (DAB) chromogen solution (DAKO) for ten minutes to develop the stain. Sections were mounted and dried on glass slides before being cover-slipped for microscopy. Adjacent sections were stained with Nissl to assist in the identification of each region of interest.

### Quantification of PV cell counts

Slides containing tissue processed for PV immunoreactivity were analyzed using a Nikon Eclipse E600 (Melville, NY) upright microscope equipped with an Insight SPOT digital camera (Diagnostic Instruments, Inc.). Photomicrographs of regions of interest including: retrosplenial cortex, somatosensory barrel fields, and HPC (including CA1, CA3, and DG sub-divisions; as sampled in Nomura et al., 1997 and Honeycutt et al., 2016), were catalogued and stored digitally for analysis by a researcher blind to treatment and age. For additional sampling and analysis parameters, see previously reported methods from our group (Honeycutt et al., 2016) and Figure1B for schematic of counting regions. To ensure precision in the quantification of PV-expressing cells within each region of interest photomicrographs were taken at either 20x magnification (CA1, CA3) or 10x magnification (cortical areas, DG). Digital photomicrographs were analyzed using ImageJ software (NIH), and the number of PV-immunoreactive cells were quantified using macros written to automate counting for each region and calculate the number of PV+ cells based on color density to isolate the most darkly stained cells, and to exclude any thresholded particles (e.g. non-cell artifacts) that did not meet criteria for inclusion (i.e. size, sphericity). A representative example of cell identification/counting using color thresholding can be seen in Figure 1C. This methodology to quantify PV as cell counts using ImageJ (as opposed to stereology) was carefully selected to address the fact that PV cells have a non-uniform distribution, particularly within the HPC and neocortex (Benes and Lange, 2001; Guillery, 2002). The sizes of the analyzed regions were as follows: HPC CA1/CA3 - 800μm^2^; HPC DG and cortical areas - 1600μm^2^. The average number of PV cells for HPC CA1 and CA3 were normalized to represent the average number of PV-positive cells per 1600μm2 for cross regional comparisons and consistency across data sets.

### Statistical analysis of PV cell count

To characterize the differences in PV+ cell counts within specified regions of interest based on treatment and time of sacrifice, the average number of PV expressing cells within each photomicrograph was calculated using ImageJ, and group means were computed. This was done across brain regions for each rat, and for each region of interest the average number of PV cells was computed from 3-6 photomicrographs per region (2-3 from each of the left and right hemispheres). Data was collapsed across hemisphere for all brain regions, as there were no significant hemispheric differences in PV count. A two-way ANOVA (Treatment (CON, KET-I, KET-D) x Age (1mo, 6mo) was conducted for each neocortical region to determine differences based on age and/or treatment and followed up with Holm-Sidak’s post-hoc analyses with corrections for multiple comparisons.

Within the HPC sub-regions (CA1, CA3, DG) averages for septotemporal regions (septal, midseptotemporal, temporal) were considered individual samples and contributed to overall averages for respective regions, such that each group’s sub-region average was comprised of 10 (DG) to 15 (CA1, CA3) data points. Differences in PV cell counts were assessed using separate two-way ANOVA analyses (Treatment (CON, KET-I, KET-D) x Age (1mo, 6mo) for each sub-region to determine whether there were changes within each group as a function of treatment and/or age. Significant effects were followed up with Holm-Sidak’s post-hoc analyses with corrections for multiple comparisons.

## Results

### Hippocampal findings

#### Age-dependent PV cell counts in CON groups

In the HPC there were significant main effects of age (*F*(1, 74) = 24.07, *p* < 0.01) and region (*F*(2, 74) = 12.59, *p* < 0.01). Specifically, we observed an overall decrease in PV cell count as a function of age between 1- and 6-month-old CON groups, such that 6-month-old rats had significantly lower PV cell counts than 1-month-old rats. This age-dependent decrease in PV was seen across all HPC sub-regions: CA1 (*p* < 0.01), CA3 (*p* < 0.01), and DG (*p* < 0.05).

#### Chronic KET administration and time of sacrifice as a function of age

In CA1 there was a significant Age x Treatment interaction (*F*(2,84) = 5.34, *p* < 0.01), and this significant interaction was also observed in CA3 (*F*(2, 84) = 5.89, *p* < 0.01) and DG (*F*(2, 54) = 9.99, *p* < 0.001). Post-hoc Holm-Sidak’s multiple comparisons of simple effects within each age group were conducted to evaluate significant interactions observed in CA1, CA3, and DG. In 1-month-old rats, chronic KET treatment significantly reduced PV cell count in both KET-I and KET-D groups compared to CON in CA1 (*p* < 0.05), CA3 (*p* < 0.05), and DG (*p* < 0.01). Notably, there was no effect of tissue collection timeline, evidenced by a lack of difference in PV count between KET-I and KET-D groups. See Figure 2 for photomicrographs of HPC regions and Figure 3 for graphical data.

**Figure 2.**
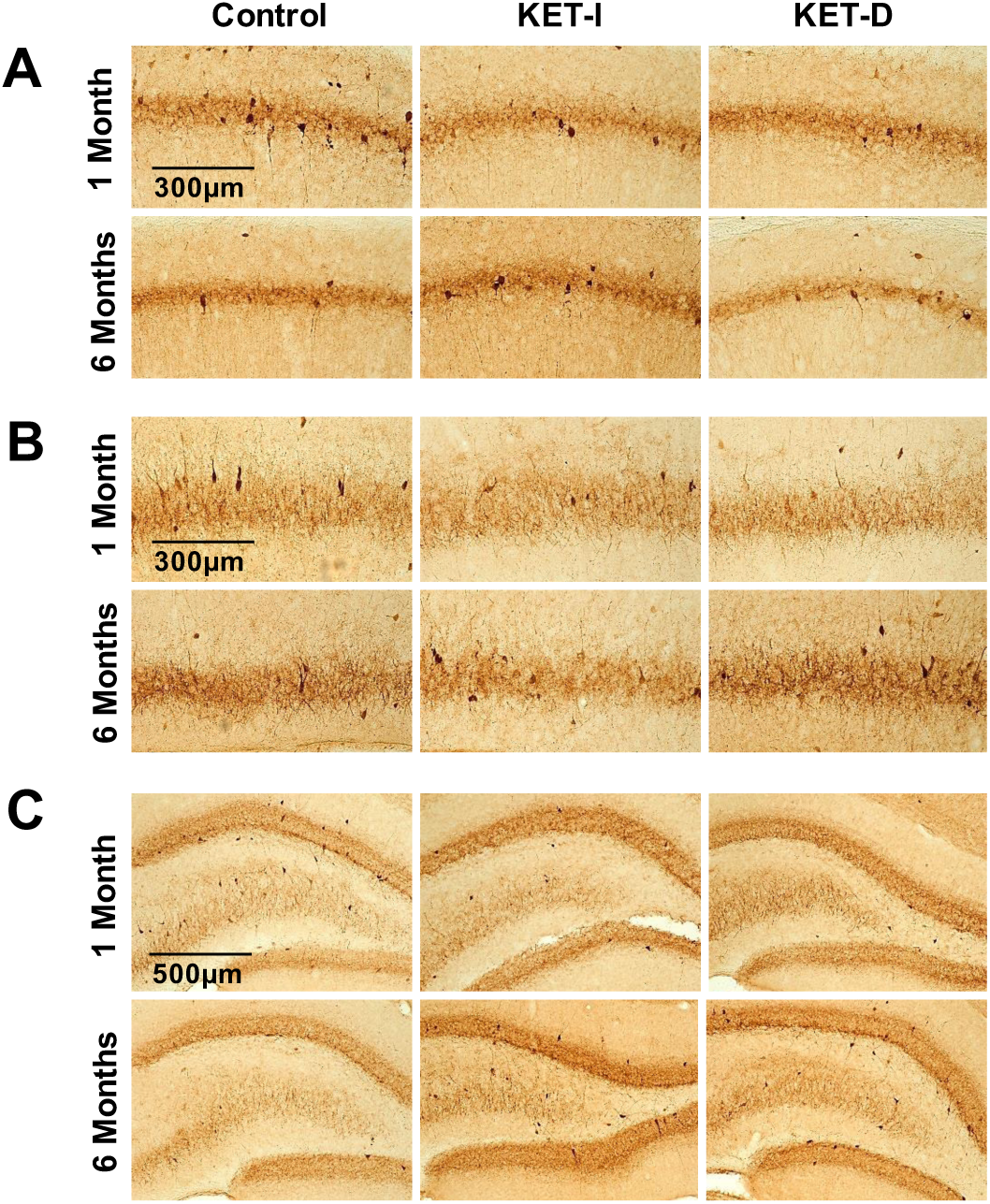
Representative photomicrographs of HPC sub-regions used for analysis in the present study: CA1 (**A**), CA3 (**B**), and DG (**C**) in 1-and 6-month CON, KET-I, and KET-D groups. There was a significant decrease in PV cell count in all sub-regions in 1-month old animals following KET treatment, while 6-month old rats showed a general increase in mean PV cell count following KET treatment (significant increase noted in DG). Time of tissue collection (KET-I v. KET-D) did not significantly impact results.

**Figure 3.**
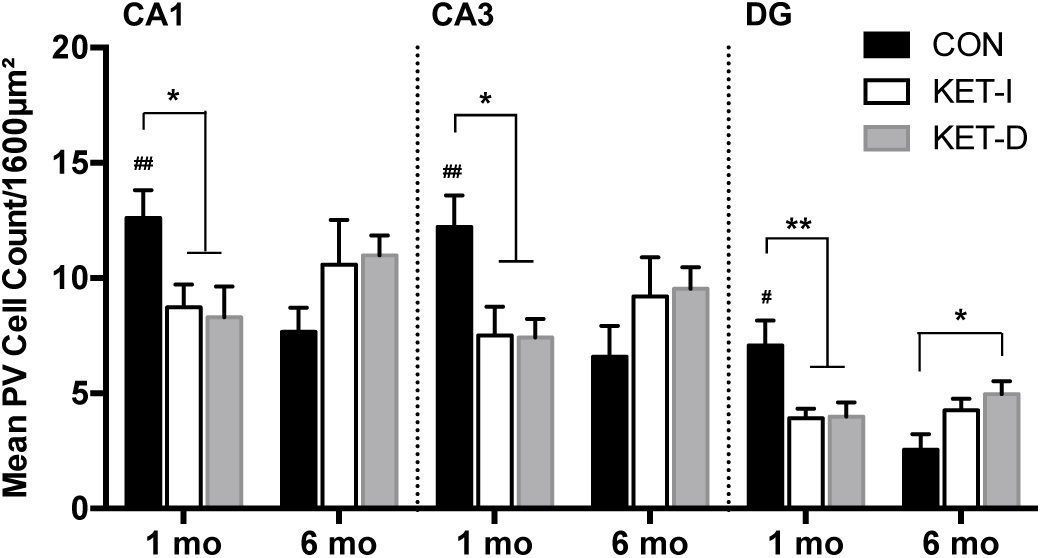
Chronic KET treatment significantly decreased mean PV cell count across all HPC sub-regions (CA1, CA3, DG) exclusively in 1-month old rats (**\****p* < 0.05; **\*\****p* < 0.01). Conversely, chronic KET treatment in 6-month old rats generally increased mean PV cell count, reaching significance within the DG. Time of tissue collection (KET-I v. KET-D) did not meaningfully impact cell count in any groups. A significant developmentally dependent decrease in mean PV cell count between 1-and 6-month old CON rats was seen across all HPC sub-regions. Significant differences between CON groups in each sub-region are indicated with a **#** (*p* < 0.05) or **##** (*p* < 0.01).

In 6-month-old rats, KET treatment generally did not significantly alter PV cell count within CA1 or CA3. Within the DG, however, post-hoc comparisons revealed a significant *increase* in PV cell count in KET-D rats compared to CON (*p* < 0.05) and a trending difference between KET-I and CON groups (*p* = 0.078). Additionally, there was an overall observable trend toward an increase in PV cell count between CON and KET-I/KET-D treated groups in both CA1 (*p* = 0.134) and CA3 (*p* = 0.196) regions. Overall, there were no significant differences based on time of tissue collection between KET-I and KET-D groups. See Figures 2 and 3.

### Neocortical findings

Within the retrosplenial cortex (Figure 4A), there was no main effect of age (*F*(1, 24) = 0.26, *p* = 0.61), or treatment (*F*(1, 24) = 0.88, *p* = 0.43). Similarly, there was no main effect of age (*F*(1, 24) = 2.78, *p* = 0.11) or treatment (*F*(1, 24) = 0.86, *p* = 0.44) in the somatosensory barrel fields(Figure 4B). In addition, neither cortical region (retrosplenial nor somatosensory) showed an interaction effect between the two factors (*p* = 0.37 and 0.63, respectively). Representative photomicrographs can be seen in Figure 4A-B.

**Figure 4.**
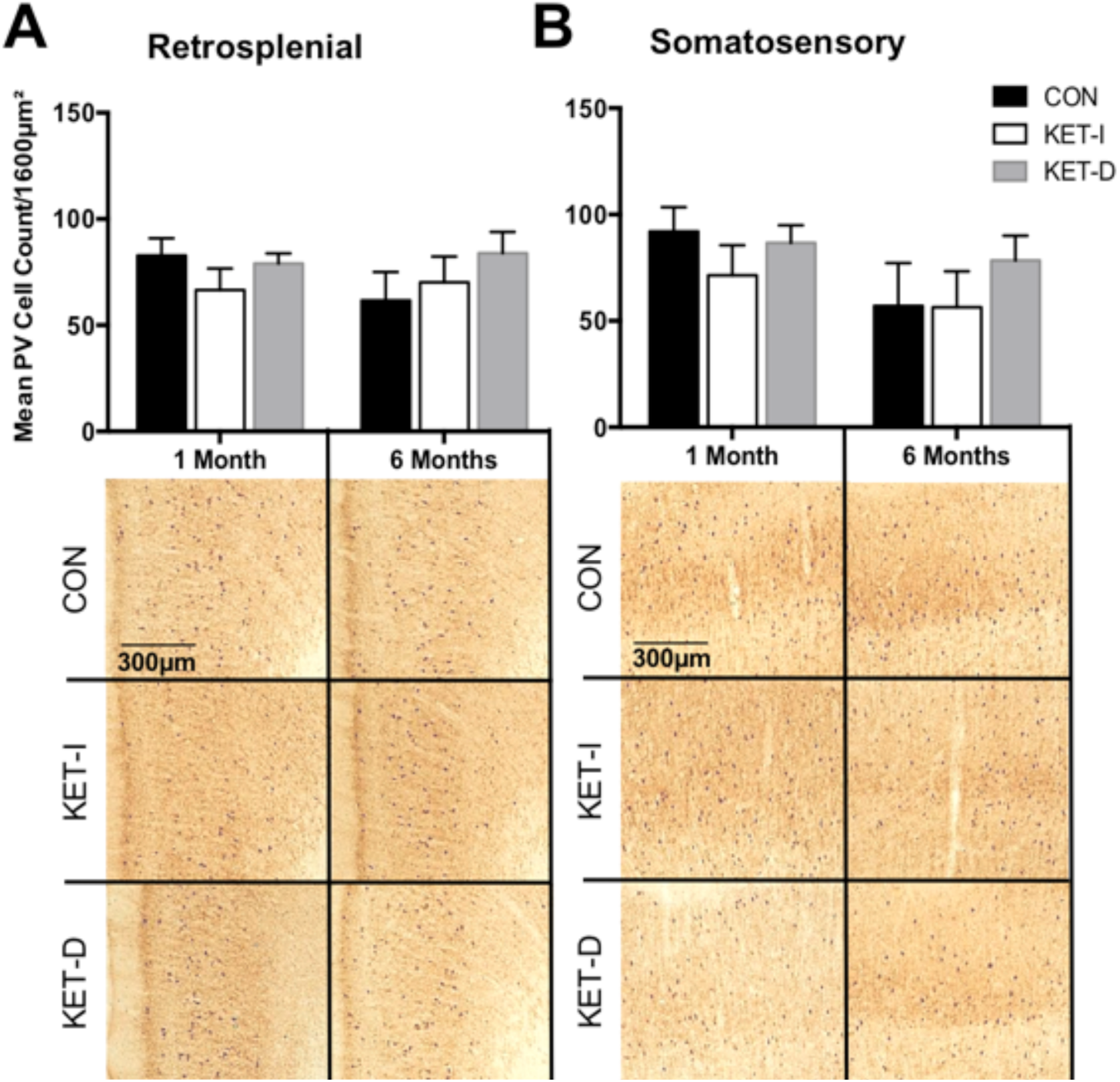
Representative photomicrographs and results of cortical analyses. Retrosplenial (**A**) and Somatosensory Barrel Field (**B**) cortices did not show changes in PV cell count as a function of age. Furthermore, none of the neocortical regions examined showed changes in PV cell count following KET administration in either KET-I or KET-D groups. Representative photomicrographs of these cortical data can be seen below the graphs.

## Discussion

The present work supports previous studies detailing a developmentally-dependent normative decrease in PV cell count within all sub-regions of the HPC (CA1, CA3, DG), but not within adjacent cortical regions in control rats (Honeycutt et al., 2016). Importantly, this work is the first to detail a novel dissociation between the effects of chronic KET administration on PV cell count based on age of treatment and anatomical region, without regard to time of tissue collection. Here, we discuss the main findings observed, including: 1) age as a significant factor in normative PV cell count within the HPC, a key brain region related to schizophrenia; and 2) the differential modulation of chronic sub-anesthetic KET administration on PV cell count as a function of developmental age.

### Age significantly impacts normative PV counts

Overall, there was a main effect of age in CON rats, such that 1-month-old rats had significantly higher PV cell counts within the HPC than 6-month-old CON rats. Notably, these findings detail a marked (up to approx. 50%) decrease in PV across all three HPC sub-regions (CA1, CA3, and DG) from 1 to 6-month CON animals, with the most dramatic age-related decrease observed in DG. This, in conjunction with prior reports of age-dependent decreases from our lab (Honeycutt et al., 2016) and others (Lolova and Davidoff, 1992; Miettinen et al., 1993; Vela et al., 2003; Lee et al., 2013), provide evidence for natural alterations in PV phenotype across development in a region-specific manner. Importantly, the present findings provide clarity to this body of work, as the rats in this study were behaviorally naïve, whereas many studies employ behavioral assays on their subjects that may alter baseline PV due to enrichment effects of training (e.g., Iuvone et al., 1996; Komitova et al., 2013). These robust changes in PV cell count, as a function of age, should be thoroughly considered, as differences in baseline counts can interfere with experimental manipulations, leading to lack of validity as well as difficulty in interpretation and generalization of results.

### Chronic KET treatment differentially impacts PV cell count as a function of age and region

The present work reveals that chronic KET administration selectively altered PV cell count within the HPC but did not significantly change the number of PV+ cells within the neocortical areas examined in either 1-or 6-month old rats, nor did time of sacrifice. Importantly, the present findings indicate a differential and opposite role of chronic KET treatment on PV cell count that is age-dependent within the HPC.

We present robust evidence for a decrease in PV cell counts within 1-month-old KET treated rats compared to age-matched CON, which is in agreement with previous work (e.g., Kittelberger et al., 2012). The reduction in PV cell count is nearly 30% in KET treated groups across all HPC sub-regions, and remains consistent regardless of time of sacrifice, indicating a long-term sustained effect of chronic KET treatment on underlying neuropathology. Conversely, we also describe an opposite effect in 6-month-old KET treated rats, such that those treated show a general increase in PV cell counts compared to their age-matched CON counterparts. The observation of a significant reduction in PV cell count in 1-vs. 6-month-old rats suggests that KET may be interacting with PV expressing neurons in different ways based on age and age-related changes in PV-localized NMDA receptors (Cull-Candy et al., 2001). This dissociation likely also speaks to the developmental switch in NMDA receptor subunit composition, which has been long theorized to be a contributing factor to the delayed onset of schizophrenia symptoms in late adolescence/early adulthood (Law et al., 2003).

In addition to its blockade of NMDA receptors, it should be noted that KET has a wide range of molecular targets, particularly within brain regions (i.e. HPC) necessary for cognitive function (Sleigh et al., 2014). Indeed, KET also interacts with HCN1 receptors (Chen et al., 2009), and nicotinic acetylcholine receptors (Scheller et al., 1996), with alterations in the function of these receptors implicated in cognitive dysfunction (e.g., Levin, 2012; Luo et al., 2015). KET administration is also linked to alterations in the release and modulatory functions of neurotransmitters such as acetylcholine (Lydic and Baghdoyan, 2002; Nelson et al., 2002) and dopamine (Kamiyama et al., 2011), and has been shown to mediate GABA function (i.e. McNally et al., 2011; Wang et al., 2017). Because PV is expressed exclusively on GABAergic neurons, it is likely that KET-induced alterations in the function of these neurons – via the aforementioned mechanisms – may differentially contribute to the changes seen in PV cells following chronic treatment. As such, additional studies are needed to further elucidate upon which of these pathways lead to measurable changes in PV cell count following KET.

The increase in PV cell count in adult rats described in the present study has not been well documented in the literature, though our findings corroborate reports of an increase in HPC PV (Sabbagh et al., 2013) and cortical PV (Abdul-Monim et al., 2007) cell counts using a similar treatment protocol. Indeed, PV has been shown to be dynamic and impacted through a variety of avenues, including behavioral and/or environmental enrichment experiences (Gomes da Silva et al., 2010; Urakawa et al., 2013; Komitova et al., 2013; Donato et al., 2014; Sun et al., 2016; Sampedro-Piquero et al., 2016), though the mechanism by which KET is having a similar age-dependent impact warrants further investigation. While it is unclear the mechanism by which PV, specifically, can be up-and/or down-regulated, evidence suggests that chronic low-dose KET administration has the capacity to increase neural proliferation and functional maturation within the HPC in adult rats (Soumier et al., 2016). This KET-induced increase in neural proliferation, paired with evidence that progenitor cells are capable of differentiating into PV+ GABAergic well into the postnatal period (Shetty and Hattiangady, 2013), may in part explain/contribute to the increased DG PV+ cells following KET treatment in 6mo rats observed here. Furthermore, based on these findings, it is reasonable to conclude that KET administration may serve to equalize PV cell count between 1-and 6-month old groups via different developmentally-dependent pathways. This caveat can be extended to previous and future work to highlight the importance of controlling for – and reporting – age of treatment and/or tissue collection to better understand the role of NMDA hypofunction on vulnerable populations of GABAergic neurons implicated in schizophrenia neuropathology. It is also notable to point out the discrepancies in NMDA antagonist dosage and treatment length in previous work, as these likely also significantly contribute to variability across studies (see Table 1). While the present work focused on changes attributable to age and time of post-treatment tissue collection, the field would benefit from future research directly comparing differences in dose and treatment length on PV outcomes.

The present work highlights the importance of considering age when evaluating changes in PV cell count, particularly within the HPC, in rodent models. Specifically, this work reinforces the importance for controlling for this developmental variable when examining KET-and, by extension, NMDA antagonist - oriented models of the neuropathology of schizophrenia. We provide novel evidence indicating that chronic sub-anesthetic KET-induced changes in PV cell count may be exclusively seen in young rats, and that the timeline of administration is likely crucial to understanding the relationship between PV and KET-induced cognitive and behavioral impairments. To our knowledge, this is the first report of a systematically described differential effect of KET as a function of age – particularly within the HPC – and provides important nuance to the literature describing KET-induced changes in PV. Future studies providing a systematic characterization of the PFC and it subregions in a similar manner would be a worthwhile and substantial expansion of the present work, as alterations in PV in the PFC have been reported in schizophrenia (i.e. Lewis et al., 2012; Dienel and Lewis, 2018). Additional investigation further detailing the impact of chronic pharmacological antagonism of the NMDA receptor on concomitant cognitive-behavioral impairment across age is warranted to understand the role of PV in regulating cognition, and whether these processes may be sensitive to aging.

## Acknowledgments

The authors would like to sincerely thank the hard work of undergraduate research assistants at the University of Connecticut who assisted in the collection of data for the present manuscript, specifically: Kevin M. Keary III and Vanessa M. Kania.

